# Epidemiological Survey of Snail Intermediate Hosts of Schistosomiasis in Zaria, Northwest Nigeria and Molluscicidal Screening of *Ficus exasperata* Extracts

**DOI:** 10.64898/2025.12.23.695224

**Authors:** Umar Aliyu Umar, Muhammad Yunus Suraj, Umar Saidu, Mahmud Ali Umar, Auwal Adamu, Mukhtar Aliyu, Khadija Anawo Anas, Abdulazeez Muhsin Al-Fadeel, Waizik Akula Naaman, Sani Ibrahim, Mohammed Auwal Ibrahim

## Abstract

Schistosomiasis is a Snail-borne parasitic disease transmitted through water contact activities in endemic areas. It is associated with serious morbidities including anaemia, bladder cancer and infertility. Vector control efforts rely on the use of synthetic molluscicide (Niclosamide) which has been attributed to environmental toxicity. Snail intermediate hosts of schistosomiasis were obtained from water bodies and channels around Zaria, morphologically identified and screened for cercarial shedding. *F. exasperata* leaves and stem bark were obtained from natural habitat in Zaria, authenticated, dried under shade, ground into powder and subjected to extraction with distilled water, ethanol and methanol. The extracts were evaluated for phytochemical constituents, DPPH and FRAP radical scavenging activities and Molluscicidal efficacy against *Bulinus spp*. Higher percentage yield (8.62%) was obtained in Methanolic extract. Phytochemical analysis revealed the presence of compounds such as alkaloids, flavonoids, saponins, cardiac glycosides and steroids. Tannins and Phenolics have higher concentration compared to other phytochemicals detected in all the extracts. *In vitro* antioxidant analysis demonstrated moderate activity, with DPPH radical scavenging activity of 62%, 60% and 47% for methanolic, aqueous and ethanolic extracts, respectively. Molluscicidal screening of the extracts revealed dose dependent toxicity against Bulinus snails, with LC50 value of 23.59ppm, 28.96ppm, 31.51ppm for ethanolic, methanolic and aqueous extracts, respectively. These findings suggest that *F. exasperata* stem bark extracts could be exploited for development of eco-friendly botanical molluscicide. Further research is needed to identify the bioactive compounds responsible for Molluscicidal efficacy, elucidate the possible mechanisms of action, evaluate the efficacy in field trials and environmental impact.

## Introduction

Schistosomiasis is a public health problem in tropical and subtropical regions of Africa, Asia, the Caribbean and South America (World Health Organization, 2022). It is one of the neglected tropical diseases (NTDs) that infects more than 230 to 250 million people annually (Alemu et al., 2018) with an estimate of 779 million people being at risk of infection (Aula et al., 2021; World Health Organization, 2022). This disease causes 280,000 deaths annually (Aula et al., 2021). it is among the most prevalent human parasitic infections that ranks only second beneath malaria on the list of parasitic diseases (De Boni et al., 2021; Mawa et al., 2021), and with sub-Saharan African countries accounting for 90% of the schistosomiasis cases globally (Carbonell et al., 2021). It is sometimes referred to as Bilharzia, Bilharziasis or snail fever caused by *Schistosoma species* such *as S. haematobium, S. mansoni, S. japonicum* which are the commonly ones that have been isolated from human’s urine and stools respectively (Diego et al., 2024).

Bulinus snails species are responsible for larval development for the human *Schistosoma* parasite, *S. haematobium, which* is the most widespread schistosome species across Africa with risk of infection in freshwater in southern and sub-Saharan Africa including the great lakes and rivers as well as smaller bodies of water (Wepnje et al., 2023). They act as intermediate hosts for *Schistosoma haematobium* and play a crucial role in the transmission of the disease (Pennance et al., 2022). Control strategies often involve the use of molluscicide (niclosamide) to target these snails and disrupt the parasite life cycle (Tumwebaze et al., 2022).

*Ficus exasperata Vahl*, also called Sandpaper tree is an important plant commonly used in plant medicine that belongs to the genus Ficus and contains about 850 species of woody, trees, shrubs, vines, epiphytes, and hemi-epiphytes (Yahaya et al., 2022).

## Methodology

### Snail Sampling, Identification and Maintenance

Snail sampling was done in six (6) different locations around Zaria environs using hand-held scooping net in the morning between 7:30 – 9:30am for a period of 1 hour in each sampling site. Manual search and handpicking were also employed. Collected snails were transported to the laboratory at the Department of Biochemistry, Ahmadu Bello University, Zaria, where they were washed, and identified using shell morphology as described by Brown and Kristensen (1993). Rain boots and hand gloves were used for precautions against infections during the snail sampling. The snails were acclimatized in the laboratory for 5-6 hours prior to the experiment. They were maintained in aerated dechlorinated water at a temperature of 22-26°C and a pH of [6.9-7.2] and fed on boiled spinach (when required), but not during experiment.

### Snail Cercarial Shedding

All the snails collected and identified to be *Bulinus* spp. (intermediate hosts of *S. haematobium*) were subsequently placed in separate glass beakers containing 20 ml of dechlorinated tap water and exposed to artificial light for 2 hours. Subsequently, the water in the respective petri dishes were examined for cercariae using standard morphological characteristics under a dissecting microscope as described by Brown (1994).

### Plant Materials Collection

Fresh leaves and stem bark of *Ficus exasperata* were harvested from Kufena rock, Zaria (11º5’21”N, 7º39’22”E) on 23rd March, 2024. The plant was identified and authenticated by a taxonomist at the herbarium in the Department of Botany, Ahmadu Bello University, Zaria, and a voucher number (ABU070563) was deposited.

### Test Organism

Bulinus species were used for the molluscicidal assay of *F. exasperate* extracts, all other species including Lymnaea and biomphalaria snails were isolated.

### Reagents

Mayer’s reagent, Wagners, Dragenforff reagent, glacial acetic acid, Ferric chloride, sulphuric acid, olive oil, Sodium hydroxide, acetic anhydride, chloroform, lead acetate, Baljet’s reagent, Follin-ciocalteu, Gallic acid, sodium carbonate, quercetin, sodium nitrite, Aluminium chloride, bromocresol green, Hydochloric acid, phosphate buffer, Atropine, sodium carbonate, vanillin, DPPH, Potassium iodide, mercury chloride, trichloro acetic acid, Potassium ferricyanide.

### Laboratory Equipments and Apparatus

The laboratory equipents used in the study include; Water bath, Vortex, PH metre, Spectrophotometre, Table top centrifuge, refrigerator, milling machine, weighing balance, test tubes and rack, conical flasks, measuring cylinder, reagent bottle and sample, funnel.

### Preparation of Plant extracts

The leaves and stem bark of *Ficus exasperata* were gently rinsed to eliminate contaminants and air-dried under shade. A laboratory mill in the department of Botany was used to grind the dried leaves and stem bark into powder and sieved with a 1mm mesh size siever to obtain fine particles. Prior to usage, the ground plant material was kept in a dessicator. Thereafter, 50 grams of *F. exasperata* powdered samples were dissolved in 500ml of methanol, ethanol, and distilled water, respectively. The macerated extracts were allowed to stand for 48hours at room temperature with intermittent shaking for maximum solubility and filtered with a Muslin cloth, and finally through Whatman filter paper (number 1). The crude extracts were obtained by evaporation of the solvents in the filtrates under reduced pressure at 40°c in a water bath.

### Qualitative Phytochemical Analysis

The obtained plant crude extracts were qualitatively analyzed for the presence of phytochemical compounds using standard chemical tests:

### Test for Alkaloids

Few drops of Mayers, Wagners or Dragendorff reagent was added to the solution of the extract. Turbidity indicates the presence of alkaloids with cream, whitish or orange precipitate respectively.

### Test for Cardiac Glycosides (Keller-Killiani Test)

A solution of glacial acetic acid (4.0 ml) with 1 drop of 2.0% FeCl_3_ mixture was mixed with the 10 ml aqueous plant extract and 1 ml H_2_SO_4_ concentrated. A brown ring formed between the layers showed the entity of cardiac steroidal glycosides.

### Test for Tannins (Ferric Chloride Test)

About 0.5 g each portion was stirred with about 10 ml of distilled water and then filtered. Few drops of 1% ferric chloride solution were added to 2 ml of the filtrate. Occurrence of a blue-black, green or blue-green precipitate indicates the presence of tannins.

### Test for Saponins (Frothing Test)

5.0 ml of distilled water was mixed with aqueous crude plant extract in a test tube and it was mixed vigorously. The frothing was mixed with few drops of olive oil and mixed vigorously and the foam appearance showed the presence of saponins.

### Test for Flavonoids (Alkaline reagent test)

2 ml of 2.0% NaOH mixture was mixed with aqueous plant crude extract; concentrated yellow color was produced, which became colorless when we added 2 drops of diluted acid to mixture. This result showed the presence of flavonoids.

### Test for Terpenoids (Liebermann Burchad test)

When added 2.0 ml of acetic acid and 2 ml of chloroform with whole aqueous plant crude extract. The mixture was then cooled and we added H_2_SO_4_ concentrated. Green color showed the entity of aglycone, steroidal part of glycosides.

### Test for Steroids (Salkowski’s test)

Concentrated H_2_SO_4_ (2 ml) was added to the whole aqueous plant crude extract. A reddish brown color formed indicate the presence of steroidal aglycone part of the glycoside.

### Test for Phenols (Lead Acetate test)

Few drops of lead acetate solution was added to the solution of the extract. A yellow colored ppt indicates presence of phenols.

### Quantitative Phytochemical Analysis

The phytochemicals detected in abundance in the extract of *F. exasperata* were thereafter quantified. Additionally, quantitative analysis involved determining the concentrations of specific compounds using established calibration curves.

### Determination of Cardiac Glycoside

Cardiac glycosides were quantitatively determined according to Solich et al, (1992) with alterations. Briefly, 1 mL solution of the extract was mixed with 10 mL freshly prepared Baljet’s reagent (95 mL of 1% picric acid + 5 mL of 10% NaOH). After an hour, the mixture was diluted with 20 mL distilled water and the absorbance was measured at 495 nm UV/VIS spectrophotometer. For preparation of the standard curve, 1 mL of different concentrations (20-100 mg/L) of securidaside was prepared. Securidaside was isolated from Securigera securidaca extract. Total glycosides from triplicate replicates were expressed as mg of securidaside per gram of dried extracts.

### Steroids

The Liebermann-Bur chard reaction method was used to detect sterols and terpenoids that give dark pink to green colour, due to the hydroxyl group reacting with acetic anhydride and H2SO4. Varying concentrations of cholesterol (10-100 μg/ml) was used for standard calibration curve, which was read spectrophotometrically at 640 nm. The concentration of steroids was expressed in milligrams/gram of the crude extract (Harborne, 1973).

### Total Triterpenoid Content

1.0 mL of the solution of the extract was mixed with vanillin-glacial acetic acid solution 1.5 mL, 5% w/v) and perchloric acid solution (5.00 mL). The sample solution was heated for 45 min at 60°C and then cooled in an ice-water bath to the ambient temperature. After the addition of glacial acetic acid (2.25 mL), the absorbance was measured at 548 nm, using a UV-visible-light spectrophotometer. Ursolic acid (20–100 ug/mL in methanol) was used as a standard. Results were expressed as milligram ursolic acid equivalents (mg ursolic acid/g extract) (Harborne, 1973).

### Determination of Total Phenolics Content (TPC)

Estimation of total phenol content was measured spectrophotometrically by Folin–Ciocalteu colorimetric method, using Gallic acid as the standard and expressing results as Gallic acid equivalent (GAE) per gram of sample. Different concentrations (0.01-0.1 mg/ml) of Gallic acid was prepared in methanol. Aliquots of 0.5 ml of the test sample and each sample of the standard solution was taken, mixed with 2 ml of Folin–Ciocalteu reagent (1:10 in deionised water) and 4 ml of saturated solution of sodium carbonate (7.5% w/v). The tubes were covered with silver foils and incubated at room temperature for 30 minutes with intermittent shaking. The absorbance was taken at 765 nm using methanol as blank. The total phenol was determined with the help of standard curve prepared from pure phenolic standard (Gallic acid) (Ainsworth and Gillespie, 2007; Alhakmani *et al*., 2013).

### Determination of Total Flavonoids Content (TFC)

The TFC was determined by aluminium chloride colorimetric assay (Zhishen *et al*., 1999). Briefly, 0.5 ml aliquots of the samples and standard solution (0.01-1.0 mg/ml) of quarcetin were added to 2 ml of distilled water and subsequently with 0.15 ml of sodium nitrite (5% NaNO_2_, w/v) solution. After 6 minutes, 0.15 ml of (10% AlCl_3_, w/v) solution was added. The solutions was allowed to stand for 6 min and after that 2 ml of sodium hydroxide (4% NaOH, w/v) solution was added to the mixture. The final volume was adjusted to 5 ml with immediate addition of distilled water, mixed thoroughly and allowed to stand for another 15 min. The absorbance of each mixture was determined at 510 nm against the same mixture. TFC was determined as mg quarcetin equivalent per gram of sample with the help of calibration curve of quarcetin. All determinations were performed in triplicate.

### Determination of Total Alkaloids Content (TAC)

TAC will be quantified by spectrophotometric method. This method is based on the reaction between alkaloid and bromocresol green (BCG). The plant extract and fractions (1 mg/ml) will be dissolved in 2 N HCl and then filtered. The pH of phosphate buffer solution will be adjusted to neutral with 0.1 N NaOH. 1 ml of this solution will be transferred to a separating funnel, and then 5 ml of BCG solution along with 5 ml of phosphate buffer will be added. The mixture will be shaken and the complex formed will be extracted with chloroform by vigorous shaking. The extract will be collected in a 10 ml volumetric flask and diluted to volume with chloroform. The absorbance of the complex in chloroform will be measured at 470 nm. TAC was determined as mg Atropine equivalent per gram of sample with the help of calibration curve of Atropine. The whole experiment will be conducted in three replicates (Shamsa *et al*., 2008; Sharief *et al*., 2014).

### Determination of Total Tannins Content (TTC)

The tannins were determined by Folin - Ciocalteu method. About 0.1 ml of the sample extract will be added to a volumetric flask (10 ml) containing 7.5 ml of distilled water and 0.5 ml of Folin –Ciocalteu phenol reagent, 1 ml of 35 % Na_2_CO_3_ solution and dilute to 10 ml with distilled water. The mixture will be shaken well and kept at room temperature for 30 min. A set of reference standard solutions of Gallic acid (20, 40, 60, 80 and 100 μg/ml) will be prepared in the same manner as described earlier. Absorbance for test and standard solutions will be measured against the blank at 725 nm with an UV/Visible spectrophotometer. The tannin content will be expressed in terms of mg of GAE /g of extract (Marinova *et al*., 2005).

### Determination of Total Saponins Content (TSC)

Total saponins will be determined according to method described by Makkar *et al*. (2007). A known quantity of freeze-dried extract will be dissolved in aqueous 50% methanol and a suitable aliquot (5 mg/ml) will be taken. Vanillin reagent (0.25 ml; 8%) will be added followed by sulphuric acid (2.5 ml; 72% v/v). The reaction mixtures will be mixed well and incubated at 60°C in a water bath for 10 min. After incubation, the reaction mixtures will be cooled on ice and absorbance at 544 nm (UV visible spectrophotometer) will be read against a blank that does not contain extract. The standard calibration curve will be obtained from suitable aliquots of diosgenin (0.5 mg/ml in 50% aqueous methanol). The total saponins concentration will be expressed as mg diosgenin equivalents (DE) per g dry weight (DW).

### *In vitro* Antioxidant Analysis

The antioxidant potential of *Ficus exasperata* extracts was assessed using two well-established assays: DPPH (2,2-diphenyl-1-picrylhydrazyl) and FRAP (Ferric Reducing Antioxidant Power).

## Results and Discussion

Table 1 presents results of sampling and cercarial shedding of snails collected from water bodies around Zaria. Both *Bulinus* and *Biomphalaria* snails were recovered, however, none of the *Bulinus* snails were found to be infected. Thirty of the Biomphalaria snails were found to be infected with Schistosome parasites, 20 snails from Tohu and10 from Basawa, respectively. Results on molluscicidal screening of the *F. exasperata* extracts were presented in Table 2 and 3. Ethanolic stem bark extract has the lowest LC_50_ of 23.59, followed by methanolic (28.96) and aqueous (31.51) stem bark extracts, respectively.

**Table 1:** Snail Population and Infection Status.

| Locations | *Bulinus spp* |  | *Biomphalaria spp* |  |
| --- | --- | --- | --- | --- |
|  | No. examined | No. infected | No. examined | No. infected |
| Basawa | 48 | 0 | 290 | **10** |
| Mil Goma | 66 | 0 | 143 | 0 |
| Kwakware manu | 63 | 0 | 92 | 0 |
| Tohu | 150 | 0 | 165 | **20** |
| Total | 327 | 0 | 690 | **30** |

**Table 2:** Molluscicidal Activity of *Ficus exasperata* Leaves and Stem Bark Extracts.

| Sample | Molluscicidal screening | Molluscicidal activities (24hrs) | Post treatment (24hrs) | Total death | Mortality rate (%) |
| --- | --- | --- | --- | --- | --- |
| Control (n=12) | 0 | 0 | 0 | 0 | 0 |
| LW (ppm) | 12.50 | 0 | 1 | 1 | 10 |
| (n = 10) | 25.00 | 0 | 0 | 0 | 0 |
|  | 50.00 | 0 | 0 | 0 | 0 |
|  | 100.00 | 0 | 0 | 0 | 0 |
| SW (ppm) | 12.50 | 0 | 1 | 1 | 8.33 |
| (n = 12) | 25.00 | 1 | 0 | 1 | 8.33 |
|  | 50.00 | 12 | 0 | 12 | 100.00 |
|  | 100.00 | 12 | 0 | 12 | 100.00 |
| LE (ppm) | 12.50 | 0 | 0 | 0 | 0 |
| (n = 12) | 25.00 | 1 | 0 | 1 | 8.33 |
|  | 50.00 | 5 | 0 | 5 | 41.67 |
|  | 100.00 | 12 | 0 | 12 | 100.00 |
| SE (ppm) | 12.50 | 3 | 0 | 3 | 25.00 |
| (n = 12) | 25.00 | 6 | 1 | 7 | 58.33 |
|  | 50.00 | 10 | 0 | 10 | 83.33 |
|  | 100.00 | 12 | 0 | 12 | 100.00 |
| LM (ppm) | 12.50 | 0 | 0 | 0 | 0 |
| (n = 10) | 25.00 | 0 | 1 | 1 | 10.00 |
|  | 50.00 | 0 | 2 | 2 | 20.00 |
|  | 100.00 | 0 | 0 | 0 | 0 |
| SM (ppm) | 12.50 | 0 | 0 | 1 | 8.33 |
| (n = 12) | 25.00 | 1 | 2 | 3 | 25.00 |
|  | 50.00 | 12 | 0 | 12 | 100.00 |
|  | 100.00 | 12 | 0 | 12 | 100.00 |

**Table 3:** Lethal Concentrations of Leaves and Stem Bark Extracts of *Ficus exasperata*.

| Conc (ppm) | SM | SE | SW | LM | LE | LW |
| --- | --- | --- | --- | --- | --- | --- |
| 12.50 | 8.33 | 25.00 | 8.33 | 0.00 | 0.00 | 10.00 |
| 25.00 | 25.00 | 58.33 | 8.33 | 10.00 | 8.33 | 0.00 |
| 50.00 | 100.00 | 83.33 | 100.00 | 20.00 | 41.67 | 0.00 |
| 100.00 | 100.00 | 100.00 | 100.00 | 0.00 | 100.00 | 0.00 |
| LC50 | 28.96 | 23.59 | 31.51 | i | 66.01 | 194 |
| LC90 | 47.91 | 52.94 | 51.56 | i | 115.50 | 235.27 |

NB: i indicate complex number, LC indicate Lethal Concentrations and ppm indicate parts per million

Table 4 presents phytochemical profile of different solvent extract of *F. exasperata* leaves and stem bark. All the phytochemicals assayed were present in all the extracts, except alkaloids which were absent in leaves extracts as well as steroids and terpenoids in aqueous leaves extract. Table 5 reveals the phytochemical composition detected in the aqueous, ethanolic and methanolic extracts of *F. exasperata* leaves and stem bark. Tannins and phenolics were detected in abundance, flavonoids and saponins present in moderate amounts while steroids and terpenoids were found in trace quantities.

**Table 4:** Phytochemical Profile of *Ficus exasperata* Leaves and Stem Bark Extracts.

| Phytochemicals | Aqueous Extract |  | Ethanolic Extract |  | Methanolic Extract |  |
| --- | --- | --- | --- | --- | --- | --- |
|  | Leaves | Stem Bark | Leaves | Stem Bark | Leaves | Stem Bark |
| Alkaloids | - | + | - | + | - | + |
| Flavonoids | + | ++ | ++ | ++ | + | + |
| Saponins | + | ++ | +++ | ++ | + | ++ |
| Tannins | ++ | +++ | +++ | +++ | ++ | +++ |
| Cardiac Glycosides | ++ | + | ++ | + | ++ | ++ |
| Steroids | - | + | + | + | + | + |
| Terpenoids | - | + | + | + | + | + |
| Phenolics | ++ | +++ | ++ | +++ | ++ | +++ |

Note: Absent (-) Trace amount (+) Moderate (++) Abundance (+++)

**Table 5:**
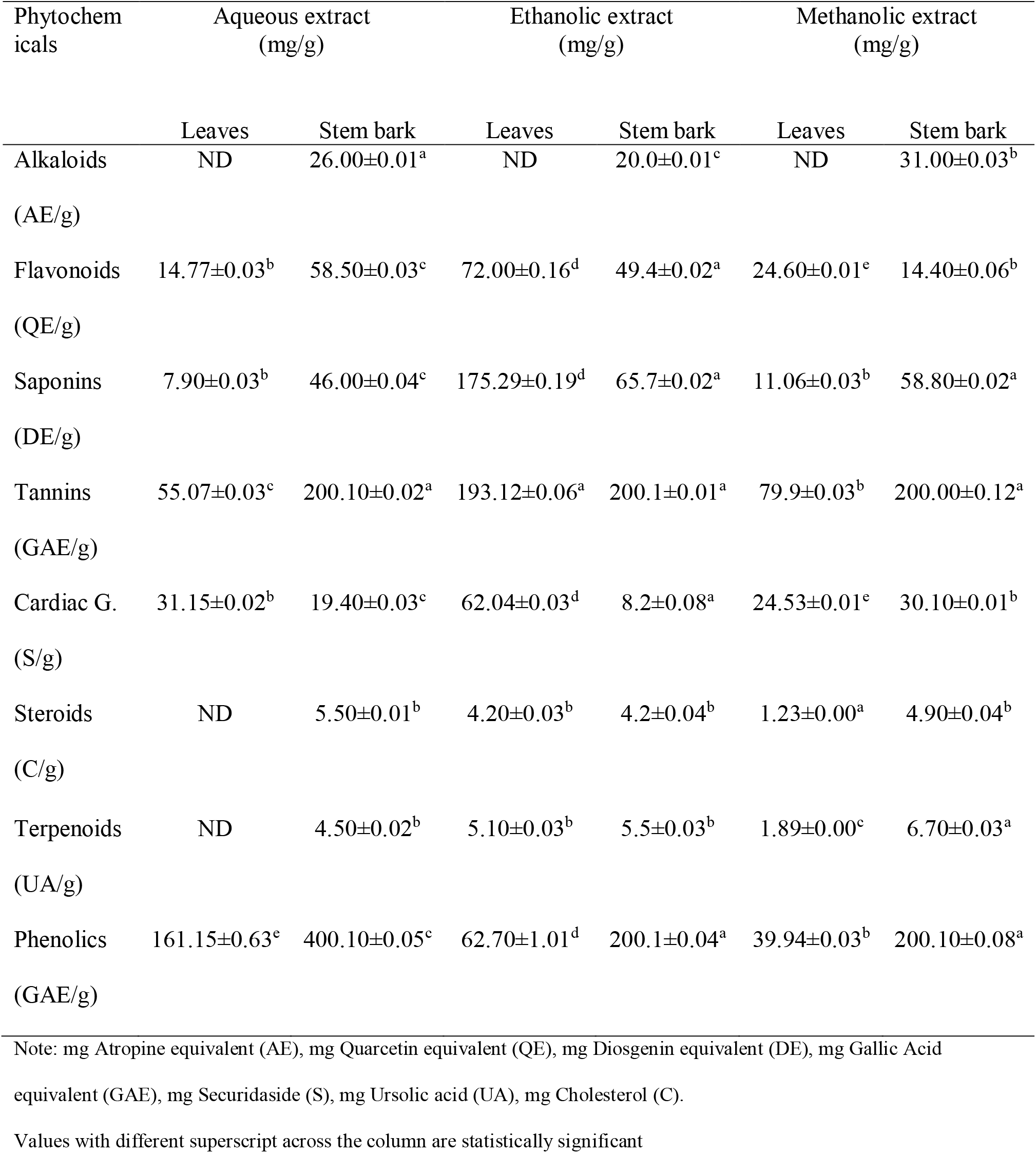
Phytochemical Composition of *Ficus exasperata* Leaves and Stem Bark Extracts.

| Phytochemicals | Aqueous extract (mg/g) |  | Ethanollic extract (mg/g) |  | Methanolic extract (mg/g) |  |
| --- | --- | --- | --- | --- | --- | --- |
|  | Leaves | Stem bark | Leaves | Stem bark | Leaves | Stem bark |
| Alkaloids<br>(AE/g) | ND | 26.00±0.01<sup>a</sup> | ND | 20.0±0.01<sup>c</sup> | ND | 31.00±0.03<sup>b</sup> |
| Flavonoids<br>(QE/g) | 14.77±0.03<sup>b</sup> | 58.50±0.03<sup>c</sup> | 72.00±0.16<sup>d</sup> | 49.4±0.02<sup>a</sup> | 24.60±0.01<sup>e</sup> | 14.40±0.06<sup>b</sup> |
| Saponins<br>(DE/g) | 7.90±0.03<sup>b</sup> | 46.00±0.04<sup>c</sup> | 175.29±0.19<sup>d</sup> | 65.7±0.02<sup>a</sup> | 11.06±0.03<sup>b</sup> | 58.80±0.02<sup>a</sup> |
| Tannins<br>(GAE/g) | 55.07±0.03<sup>c</sup> | 200.10±0.02<sup>a</sup> | 193.12±0.06<sup>a</sup> | 200.1±0.01<sup>a</sup> | 79.9±0.03<sup>b</sup> | 200.00±0.12<sup>a</sup> |
| Cardiac G.<br>(S/g) | 31.15±0.02<sup>b</sup> | 19.40±0.03<sup>c</sup> | 62.04±0.03<sup>d</sup> | 8.2±0.08<sup>a</sup> | 24.53±0.01<sup>e</sup> | 30.10±0.01<sup>b</sup> |
| Steroids<br>(C/g) | ND | 5.50±0.01<sup>b</sup> | 4.20±0.03<sup>b</sup> | 4.2±0.04<sup>b</sup> | 1.23±0.00<sup>a</sup> | 4.90±0.04<sup>b</sup> |
| Terpenoids<br>(UA/g) | ND | 4.50±0.02<sup>b</sup> | 5.10±0.03<sup>b</sup> | 5.5±0.03<sup>b</sup> | 1.89±0.00<sup>c</sup> | 6.70±0.03<sup>a</sup> |
| Phenolics<br>(GAE/g) | 161.15±0.63<sup>e</sup> | 400.10±0.05<sup>c</sup> | 62.70±1.01<sup>d</sup> | 200.1±0.04<sup>a</sup> | 39.94±0.03<sup>b</sup> | 200.10±0.08<sup>a</sup> |

Note: mg Atropine equivalent (AE), mg Quarcetin equivalent (QE), mg Diosgenin equivalent (DE), mg Gallic Acid equivalent (GAE), mg Securidaside (S), mg Ursolic acid (UA), mg Cholesterol (C).
Values with different superscript across the column are statistically significant

Figure 1 shows the DPPH radical scavenging activity of the F. exasperata extracts in comparison to ascorbic acid. Aqueous and methanolic stem bark extracts exhibited higher antioxidant activity, but are relatively low compared to ascorbic acid. Results of ferric reducing antioxidant power (FRAP) analysis of the extracts were presented in Figure 2. All the extracts exhibit low reducing power compared to the standard control (ascorbic acid), but stem bark extracts have slightly higher activity than leaves extracts.

**Figure 1.**
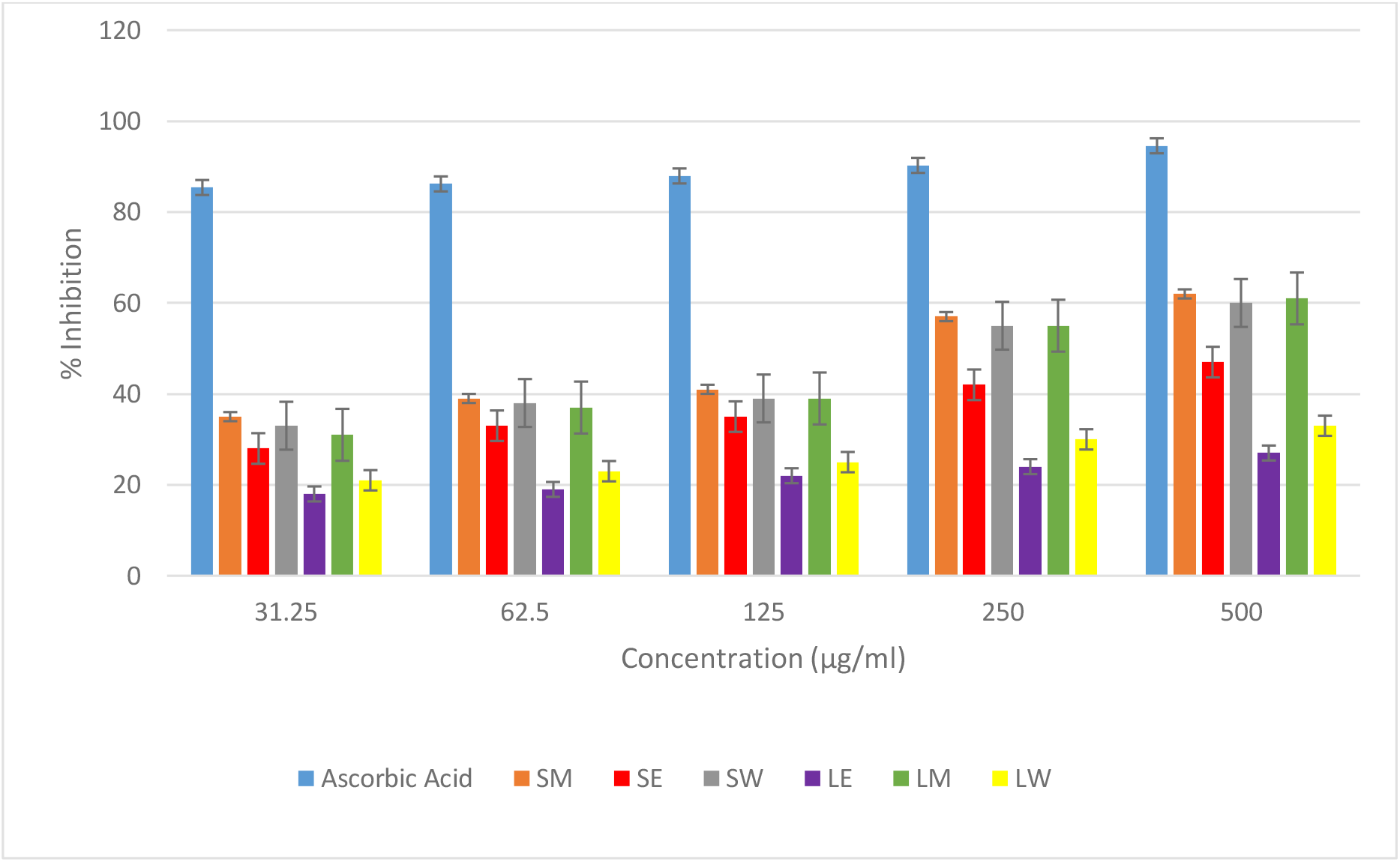
DPPH Radical Scavenging Activity of *Ficus exasperata* Leaves and Stem Bark Extracts

**Figure 2.**
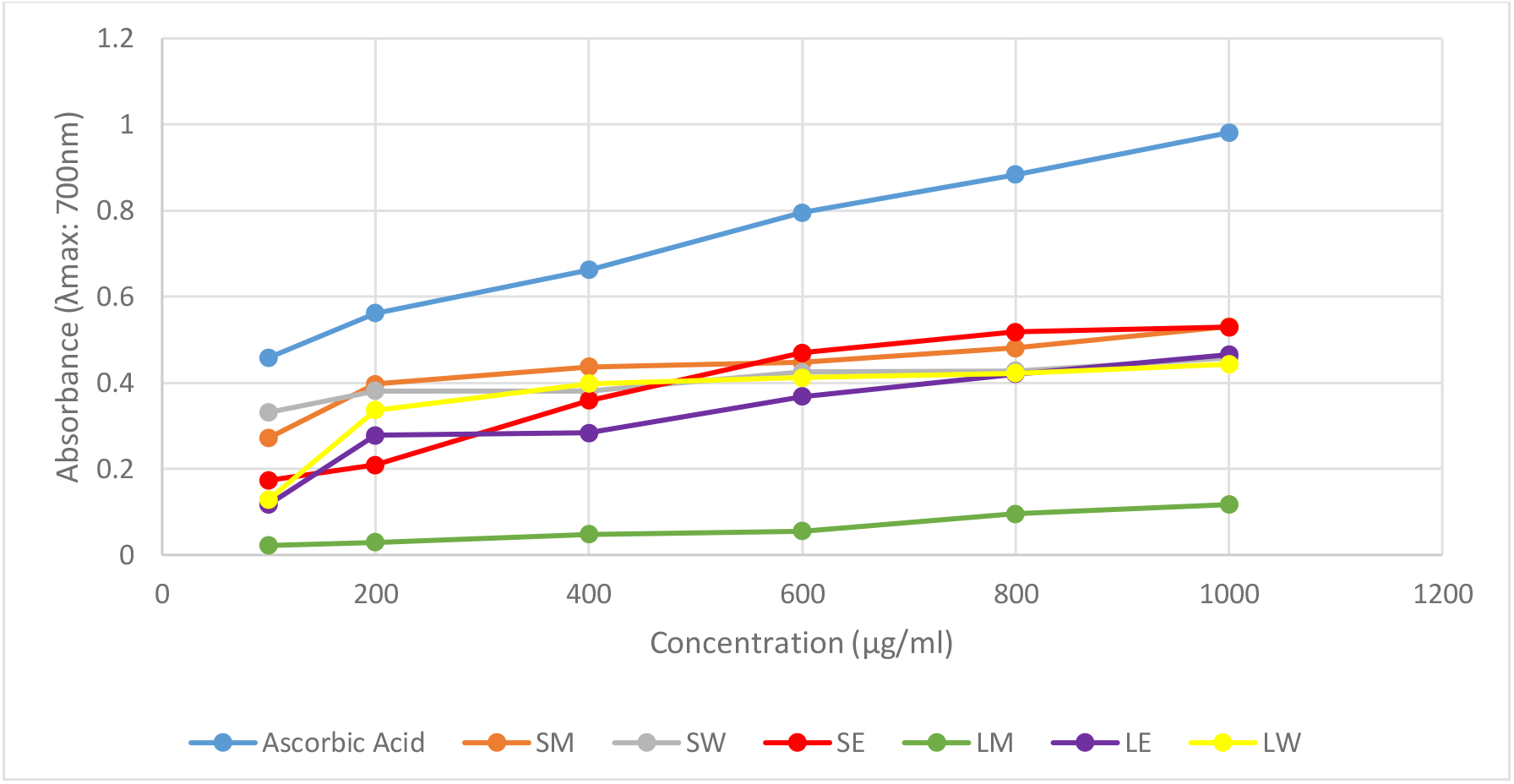
Ferric Reducing Antioxidant Power (FRAP) Activity of *Ficus exasperata* Leaves and Stem Bark Extracts

